# Toxicity of 4-(Methylnitrosamino)-1-(3-pyridyl)-1-butanone (NKK) in early development: a wide-scope metabolomics assay in zebrafish embryos

**DOI:** 10.1101/2021.06.07.447361

**Authors:** Carla Merino, Marta Casado, Benjamí Piña, Maria Vinaixa, Noelia Ramírez

## Abstract

The tobacco-specific nitrosamine 4-(Methylnitrosamino)-1-(3-pyridyl)-1-butanone (NNK) is a carcinogenic and ubiquitous environmental pollutant which carcinogenic and cytotoxic activity has been thoroughly investigated in murine models and human tissues. However, its potential deleterious effects on vertebrate early development are yet poorly understood. In this work, we characterized the impact of NNK exposure during early developmental stages of zebrafish embryos, a known alternative model for mammalian toxicity studies. Embryos exposed to different NNK concentrations were monitored for lethality and for the appearance of malformations during the first five days after fertilization. LC/MS-based untargeted metabolomics was subsequently performed for a wide-scope assay of NNK-related metabolic alterations. Our results revealed the presence of not only the parental compound, but also of two NKK known metabolites, 4-Hydroxy-4-(3-pyridyl)-butyric acid (HPBA) and 4-(Methylnitrosamino)-1-(3-pyridyl-N-oxide)-1-butanol (NNAL-N-oxide) in exposed embryos likely resulting from active CYP450-mediated α-hydroxylation and NNK detoxification pathways, respectively. This was paralleled by a disruption in purine and pyrimidine metabolisms and the activation of the base excision repair pathway. Our results confirm NNK as a harmful embryonic agent and demonstrate zebrafish embryos to be a suitable early development model to monitor NNK toxicity.

## 1. Introduction

The World Health Organization (WHO) recognizes exposure to tobacco smoke toxicants as a major environmental health issue affecting both smokers and passive smokers such as young children (World Health Organization, 2017). Almost half of children regularly breathe air polluted with tobacco smoke in public places and 65,000 die each year from illnesses attributable to secondhand smoke (SHS) which has been related to pregnancy complications and low birth weight (World Health Organization, 2020), as well as, to respiratory diseases, neurodevelopmental alterations, and increased risk of sudden infant death syndrome (U.S. Department of Health and Human Services et al., 2014). Beyond SHS, the so-called thirdhand smoke (THS) is considered as another form of passive exposure to residues of tobacco smoke that deposits, ages and remains in fabrics, surfaces and settle dust particles long after long after smoking has ceased (Matt et al., 2011).

The tobacco specific nitrosamine 4-methylnitrosamino-1-(3-pyridyl)-1-butanone (NNK) is one of the most abundant and strongest carcinogens in SHS and THS (Hecht and Hoffmann, 1988; Jacob et al., 2017; Schick and Glantz, 2007), being classified as Group 1 carcinogenic in humans by the International Agency for Research on Cancer (IARC) (International Agency for Reasearch on Cancer, 2007). NNK is formed upon the oxidation of residual nicotine from tobacco smoke with nitrous acid, ozone, and other atmospheric oxidants (Sleiman et al., 2010). NNK has been ubiquitously detected in settled dust, even in some smoke-free homes (Ramírez et al., 2014), outdoor airborne particulate matter (Farren et al., 2015), waste waters (Lai et al., 2018) and river waters (Wu et al., 2012) expanding NNK exposure to non-smoking population such as children. Relevant NNK toxicity studies have traditionally focused on its associated carcinogenicity in adult animal models. Thus, NNK has been evaluated *in vitro* in cultured explants and epithelial cells of human buccal mucosa, and *in vivo* murine models where it has demonstrated to cause tumours in nasal cavities, trachea, lung, liver, stomach, and skin (Hecht, 1998). DNA adduct formation drives these malignant transformations, a process triggered by NNK metabolic activation occurring via P450-mediated carbonyl reduction, pyridine oxidation, α-hydroxylation of the carbons adjacent to the N-nitroso group, denitrosation, and ADP adduct formation (Balbo et al., 2014). However, NNK toxicity effects during early life remain less studied even though children are considered particularly vulnerable and their exposure dose has been estimated higher than among adults (Wei et al., 2016). Some perinatal toxicity assays in murine models have demonstrated NNK to be a weak (transplacentally in mice) to moderate (neonatal mice and transplacentally in hamsters) perinatal carcinogen crossing the placental barrier and causing tumours in the offspring of exposed pregnant rodents (Anderson et al., 1991, 1989; Correa et al., 1990; Rossignol et al., 1989). NNK has also been shown to be a weak embryotoxic and teratogenic agent in mice models (Winn et al., 1998). However, mechanisms underlying such NNK toxic effects during early development remain yet to be fully elucidated.

In this context, the use of alternative animal models and the adoption of systems-wide approaches to assay toxicity has been put forward to meet the twenty-first century guidelines on toxicity testing (Hartung, 2009; National Research Council, 2007). Zebrafish *(Danio Rerio)* has emerged as an excellent vertebrate alternative model granting low-cost and high-throughput toxicological assays during developmental biology and embryogenesis. The developing zebrafish embryo has been suggested as a relevant model to screen for potential teratogens due to genetic conservation with mammals, experimental advantages over higher vertebrates and external development of transparent embryos among others (Jarque et al., 2020; Lieschke and Currie, 2007; Sipes et al., 2011). Additionally, metabolomics, that is the comprehensive measurement of metabolites in a biological specimen, can provide a direct and functional readout of an organism phenotype allowing a deeper understanding of the mode(s) and mechanism(s) of action of toxicity in human and environmental health (Ramirez et al., 2013). The goal of this study is twofold: to unveil the potential metabolic disruptions caused by NNK exposure at early developmental stages using a systemic toxicity assay based on a wide-scope metabolism interrogation through mass spectrometry-based untargeted metabolomics; and to assess whether zebrafish embryos can be used as an alternative model for NNK toxicity testing in early development.

## 2. Materials and methods

### 2.1. Chemicals and reagents

4-(Methylnitrosamino)-1-(3-pyridyl)-1-butanone (NNK) was obtained from LGC-Dr Ehrenstorfer (LGC Standards, Barcelona, Spain) and 4-Hydroxy-4-(3-pyridyl)-butyric acid (HPBA) was purchased from Toronto Research Chemicals (TRC, Ontario, Canada). The metabolites hypoxanthine, cytidine monophosphate (CMP), guanosine monophosphate (GMP), S-adenosylmethionine (SAM), and uridine monophosphate (UMP) were obtained from Sigma-Aldrich (Sigma-Aldrich, St. Louis, MO, USA). Internal standard (IS) solution was prepared with 1 mg mL^-1^ of succinic-d4 acid and myristic-d27 acid in methanol also from Sigma-Aldrich. All solvents used in the preparation of solutions were LC-MS grade. Fish water was prepared with reverse-osmosis purified water containing 90 μg mL^-1^ of Instant Ocean^®^, 0.58 mM CaSO4o2H2O (Aquarium Systems, Sarrebourg, France).

### 2.2. Zebrafish maintenance and embryo collection

Wildtype zebrafish (*Danio rerio*) embryos were obtained by natural mating of 3 males and 5 females on 4 L mesh-bottom breeding tanks. At 2 hours post-fertilization (hpf), viable fertilized eggs were collected, rinsed with fish water, and kept in clean fish water under standard conditions (28.5°C and 12 h Light:12 h Dark photoperiod) until the day of exposure. Embryos were incubated under the same conditions during all the experiment. All procedures were conducted in accordance with the institutional guidelines under a license from the local government (DAMM 7669, 7964) and were approved by the Institutional Animal Care and Use Committees at the Research and Development Centre of the Spanish Research Council, CID-CSIC.

### 2.3. NNK exposure setup

Six well-plates with 10 zebrafish embryos each were incubated from 48 hpf to 120 hpf in 5 mL of embryo water containing 50, 100, 150, 200 μM of NNK. NNK exposure concentration range was based on those previously tested in *in vitro* assays (Cheng et al., 2015; Hang et al., 2013; Weng et al., 2018; Winn et al., 1998) and those described for murine models exposed to TSNAs through drinking water (Balbo et al., 2014, 2013). Non-exposed controls were run in parallel. Ten replicates were performed for each incubation condition and media containing freshly prepared NNK solutions was renewed daily to ensure NNK continuous exposure. Anatomical development of embryos was followed daily under stereomicroscope Nikon SMZ1500 equipped with a Nikon digital Sight DS-Ri1 camera. Embryos were monitored and classified into three categories according to their phenotype: normal phenotype, severe phenotype (no-hatched embryos, oedema, and other morphological anomalies) and death embryos at 72, 96 and 120 hpf. All the parameters assessed to classify embryo phenotypes are listed in Table S1 of the Supplementary information. At 120 hpf, larvae exposed to the same NNK concentrations were randomized and the resulting pool was split into five different replicates containing 17-20 larvae each. These were rinsed with egg water (60 μg mL^-1^ of Instant Ocean^®^ sea salts in ultrapure water), snap-frozen in dry ice in 2 mL cryogenic tubes and stored at −80°C until sample extraction for metabolomics analysis.

### 2.3. Metabolite extraction

The extraction method was adapted from Raldúa D. *et al* (Raldúa et al., 2020). In brief, zebrafish larvae were homogenized in 500 μL of pre-chilled CHCl3:MeOH (2:1, v/v) and 20 μL of IS solution, with two 5 mm stainless beads using a TissueLyser (Qiagen, CA, USA) at 50 Hz for 4 min. Resulting homogenates were transferred to a 2 mL cryogenic tube, mixed with 117 μL of pre-chilled Milli-Q water and shacked for 20 min. After centrifugation (14680 rpm, 4 °C, 20 min), 100 μL of the upper aqueous fraction were transferred into amber chromatographic vials for further LC-MS analysis. Quality control samples (QCs) were prepared by pooling equal volumes of each extract.

### 2.4. LC-MS and MS/MS analysis

Untargeted metabolomics analyses were performed by injecting 3 μL of aqueous extracts in a 1290 UHPLC system coupled to a 6550 quadrupole time of flight (QTOF) mass spectrometer, both from Agilent Technologies (Palo Alto, CA, USA), operated in positive electrospray ionization mode (ESI+). LC-MS conditions were adapted from Torres S. *et al* (Torres et al., 2021). Briefly, chromatographic separation was conducted on a Luna^®^ Omega Polar C_18_ column (1.6μm, 150× 2.1mm, 100 Å, phenomenex, CA, USA) at a flow rate of 0.450 mL min^-1^. Column temperature was set to 30 °C. The mobile phases were 0.1% formic in Milli-Q water (A), and 0.1% formic in acetonitrile (B). The linear gradient elution started at 100% of A (time 0-2 min), 100% of A to 100% of B (2-9 min), 100% of B (9-10 min) and finished at 100% of A (10-12 min) followed by 4 minutes post-run time. ESI conditions were gas temperature, 150°C; drying gas, 11 L min^-1^; nebulizer, 30 psig; fragmentor, 120 V; and skimmer, 65 V. Mass spectra were acquired over the m/z range 50-1000 Da at 3 spectra s^-1^. For identification purposes, targeted MS/MS towards protonated ionic species from relevant features were performed under the same chromatographic conditions at fixed collision energies of 10, 20, 30 and 40 V. The instrument was set to acquire the selected precursor ions over the m/z range 40-1000 Da with a narrow isolation window width of 1.3 m/z.

### 2.5 Data processing and statistical analysis

Differences in teratogenic anomalies were evaluated using Z-test of proportions. LC-MS raw data was converted into mzXML format via MSconvert software from Proteowizard (Holman et al., 2014) and then processed using the XCMS software (version 3.8.2) to detect and align features (Colin A. Smith et al., 2006). An intensity threshold above 5,000 counts and a relative standard deviation higher than in QCs were used as criteria to retain features for downstream statistical analysis. Feature matrix intensities were normalized using Probabilistic Quotient Normalization (PQN) to account for sample dilution. Samples were normalized using the number of zebrafish embryos before entering a principal component analysis (PCA) for initial inspection of different exposure levels groups. One-way ANOVA with false discovery rate (FDR) control was used to study differences across exposure levels (p<0.05 were considered for significance in all statistical test). Heatmap with hierarchical clustering of the identified metabolites was performed using Pheatmap package version 1.0.12 (Kolde, 2012). Figures were created using ggplot2 version 3.3.0 (Wickham, 2011). Data processing and analysis was conducted in R version 3.6.3 (R Core Team, 2017)

### 2.6. Metabolite identification and pathway analysis

For metabolite identification purposes, exact masses associated to those metabolic features found to vary according to NNK exposure concentrations were searched against the HMDB (Wishart et al., 2018) considering the protonated adduct and a mass error within 5 ppm. Additionally, these exact masses were also searched against an in-house compiled list of previously reported NNK metabolites (Dator et al., 2018; Hecht, 1998) considering protonated adducts within 5 ppm mass error (Table S2). Identification of those metabolites was performed by matching their experimental MS/MS spectra with the MS/MS of NNK metabolites pure chemical standards— level 1 of confirmation according to Schymanski *et al.* (Schymanski et al., 2014) —. When pure chemical standards were not available, metabolic annotation (Nash and Dunn, 2019) was performed by comparing our experimental MS/MS spectra with experimental MS/MS spectra available in METLIN (Smith et al., 2005) and MoNA (“MoNA: MassBank of North America”, 2021) databases, or with MS/MS data from the bibliography (Dator et al., 2018) (level 2 of confirmation). Identifications were used to feed a graphical network-based pathway enrichment analysis via FELLA package version 1.10.0 (Picart-Armada et al., 2018) using *Danio rerio* as a background organism. Diffusion analysis method was performed with the top 150 z-score to prioritise affected KEGG pathways upon NNK exposure (p-score < 0.05 were considered for significance).

## 3. Results and discussion

### 3.1. NNK embryo toxicity assays

Exposure to NKK induced zebrafish embryo toxicity in a concentration-dependent manner (Figure 1). Based on lethality data, 200 μM was identified as the lowest-observed-adverse-effect level (LOAEL) yet increased abnormalities (spine cord modifications, development of oedema, and unhatched embryos) were observed in embryos exposed to concentrations ranging from 50 to 100 μM NKK (Figure 1). This suggests a weak NNK teratogenic and embryotoxic effect at the range of concentrations tested in our experiments. Our observations are in line with previous reports in mice, demonstrating a weak NNK-induced teratogenic effect in the progeny of CD1 pregnant mice and in an *in vitro* assay using mice embryos exposed to 10 and 100 μM NNK (Winn et al., 1998).

**Figure 1.**
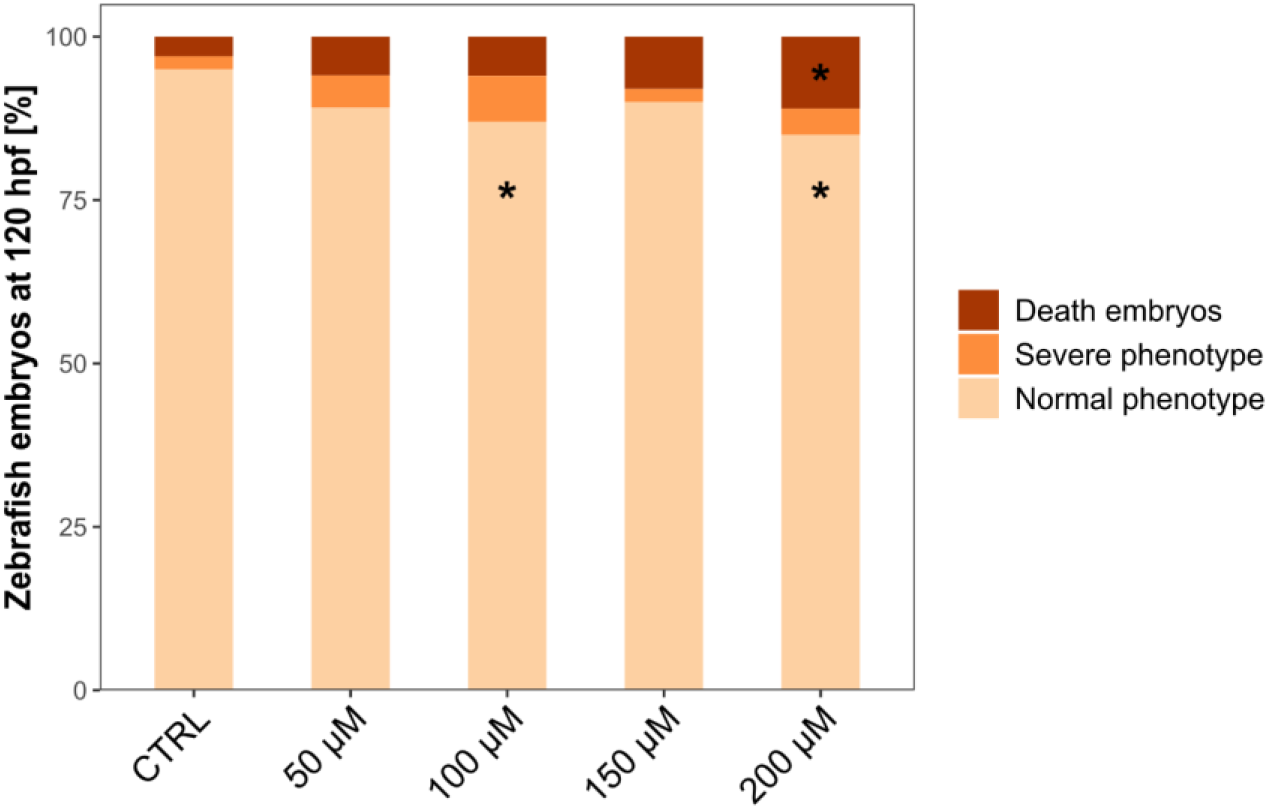
Phenotypic observations of zebrafish embryos exposed to concentrations of NNK ranging from 50 to 200 μM at 120 hpf. Observations were based on normal phenotype, severe phenotype (no-hatched embryos, oedema, and abnormalities in morphology) and death embryos. Asterisks denote statistical significance according to Z-Test compared to control group; * p-value < 0.05.

### 3.2 NNK metabolism suspect screening

To assess NNK specific metabolism in zebrafish embryos a suspect screening analysis for the 23 NNK metabolites was performed with three out of all screened metabolites identified in our samples (Table S2) namely NNK, 4-Hydroxy-4-(3-pyridyl)-butyric acid (HPBA) and 4-(Methylnitrosamino)-1-(3-pyridyl-N-oxide)-1-butanol (NNAL-*N*-oxide). Both NNK and HPBA were shown to accumulate in zebrafish embryos in a NNK concentration-dependent manner while NNAL-N-oxide reached a plateau at 100 μM (Figure 2). The accumulation of these three compounds in zebrafish embryos extracts indicate the absorption and metabolization of NNK by our zebrafish model. NNK absorption and metabolisation has been previously demonstrated in human cells, tissues from murine foetuses and in adult murine models (Hecht, 1998; Jalas et al., 2005; Rossignol et al., 1989). However, this is the first evidence demonstrating NNK to be metabolized by a zebrafish model at early developmental stages. Our results showed HPBA to be significantly increased in all NNK exposed groups respective to control group. In humans (Wang et al., 2019) and rodent models (Schrader et al., 1998), HPBA is a product of NNK metabolic activation leading to NNAL formation followed by CYP450-mediated α-hydroxylation of its carbon adjacent to the nitrosamino group (Smith et al., 1992). Of note, it has been recently demonstrated that various drug-metabolizing CYPs are expressed in zebrafish embryos and larvae and that their metabolic activity resembles to those of human CYP isoforms (Nawaji et al., 2020). This CYP450-mediated α-hydroxylation can lead to the formation of reactive intermediates that react with DNA forming methyl (i.e., O^6^-mdG, 7-mdG), pyridyloxobutyl (POB, O^6^-POB-dG) or/and pyridylhydroxybutyl (PHB) -DNA adducts which are critical in carcinogenesis (Balbo et al., 2014; Hang, 2010; Hecht et al., 2016; Hecht, 1999) (Figure 2). Our results also indicate that our zebrafish embryo model is undergoing a NNK detoxification process as suggested by the formation of NNAL-*N*-oxide which is the product of the pyridine-*N*-oxidation reaction of NNAL mediated by P450 enzymes (CYP450) (Carmella et al., 1997; Perez-Paramo et al., 2019).

**Figure 2.**
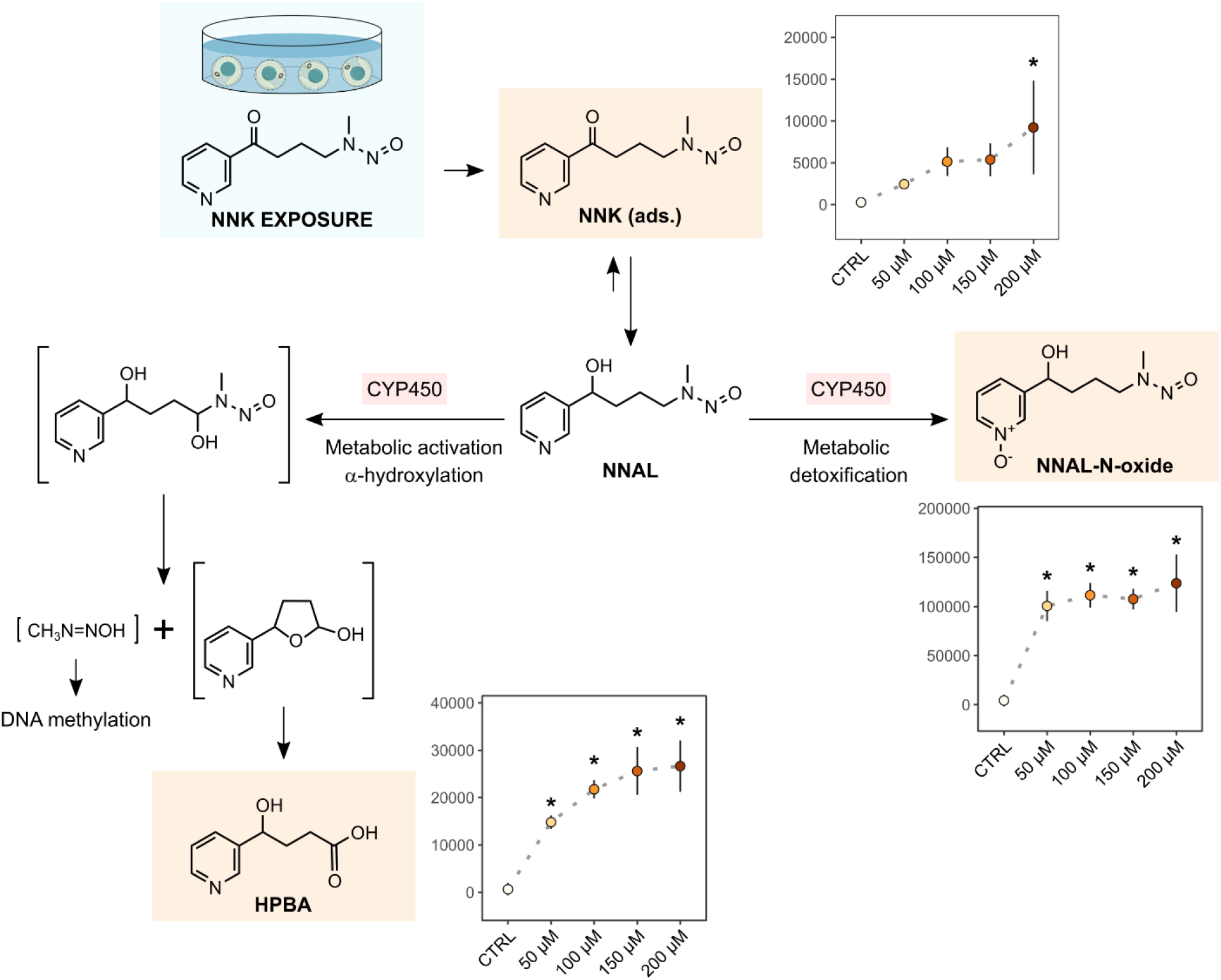
Trends of NNK derived metabolites levels identified in exposed groups. Each point represents the mean intensities per group and error bar represents the standard deviation. Asterisks denote statistical significance according to multiple group comparison of ANOVA (only significative results of exposed groups vs control are illustrated); * p-value < 0.05. n = 5 samples. NNK: 4-(methylnitrosamino)-1-(3-pyridyl)-1-butanone; NNAL-*N*-Oxide: 4-(Methylnitrosamino)-1-(3-pyridyl-N-oxide)-1-butanol; HPBA: 4-Hydroxy-4-(3-pyridyl)-butyric acid. NNK metabolism adapted from Hecht S. (Hecht, 1998).

### 3.3 NNK-related metabolic disruptions

Metabolic disruptions occurring upon NNK exposure during early development were assayed using an untargeted metabolomics approach and its results are summarized in Figure 3. was performed. Figure 3A displays PC1 vs PC2 scores plot of PCA analysis performed on the filtered and normalized metabolic features revealing a NNK concentration-dependent clustering trend along PC1 gathering 25% of explained variance. An overlap of those embryo replicates exposed to the two highest NNK concentrations (150 and 200 μM) is also evidenced, indicating similarities in their metabolic response. Metabolic features accounting for significant differences in a one-variable-at a time 1-way ANOVA comparison were used for MS/MS identification. The identity of 21 metabolites were confirmed (Table 1) including amino acids such as methionine, dipeptides, fatty acid esters, nucleobases—such as guanine and uracil—, nucleosides—such as guanosine, inosine, and uridine—, and nucleotides—such as CMP, GMP and UMP—. A heatmap representation of intensity levels for these 21 metabolites is displayed in Figure 3B, showing three clusters corresponding to the non-exposed, to the lowest and to the highest NNK exposure concentrations. The 21 identified metabolites were used as input to network-based enrichment analysis (Figure 3C) showing that three main pathways were influenced by the exposure to NNK in early development: purine and pyridine metabolisms, and base excision repair (BER) pathway. Metabolites belonging to purine metabolism, such as hypoxanthine, inosine, guanine, and guanosine, followed a decreasing trend as the concentration of NNK exposure increased (Figure 3B). In contrast, intensities of pyrimidines such as uracil and uridine showed a sharp increase at lowest exposure concentration (50 μM) and a progressive decrease with increased NNK concentrations (Figure 3B). The RNA monomers CMP, GMP and UMP followed this same trend, maintaining significantly increased concentrations respective to controls and peaking at 50 μM (Figure 4). Increased ribonucleotide biosynthesis has been linked to carcinogenic processes, which require elevated rates of RNA synthesis (Bywater et al., 2013; Villicaña et al., 2014). On the other hand, our results suggest a significant alteration of BER pathway activity upon NNK exposure, an effect not observed in mice models (Gupta et al., 2013). BER pathway is one of the primary repair mechanisms for the removal of small DNA lesions such as alkylated, oxidized, and deaminated bases from endogenous sources or environmental carcinogens, i.e., non-bulky adducts such as those produced by NNK (Hang, 2010; Peterson, 2010).

**Figure 3.**
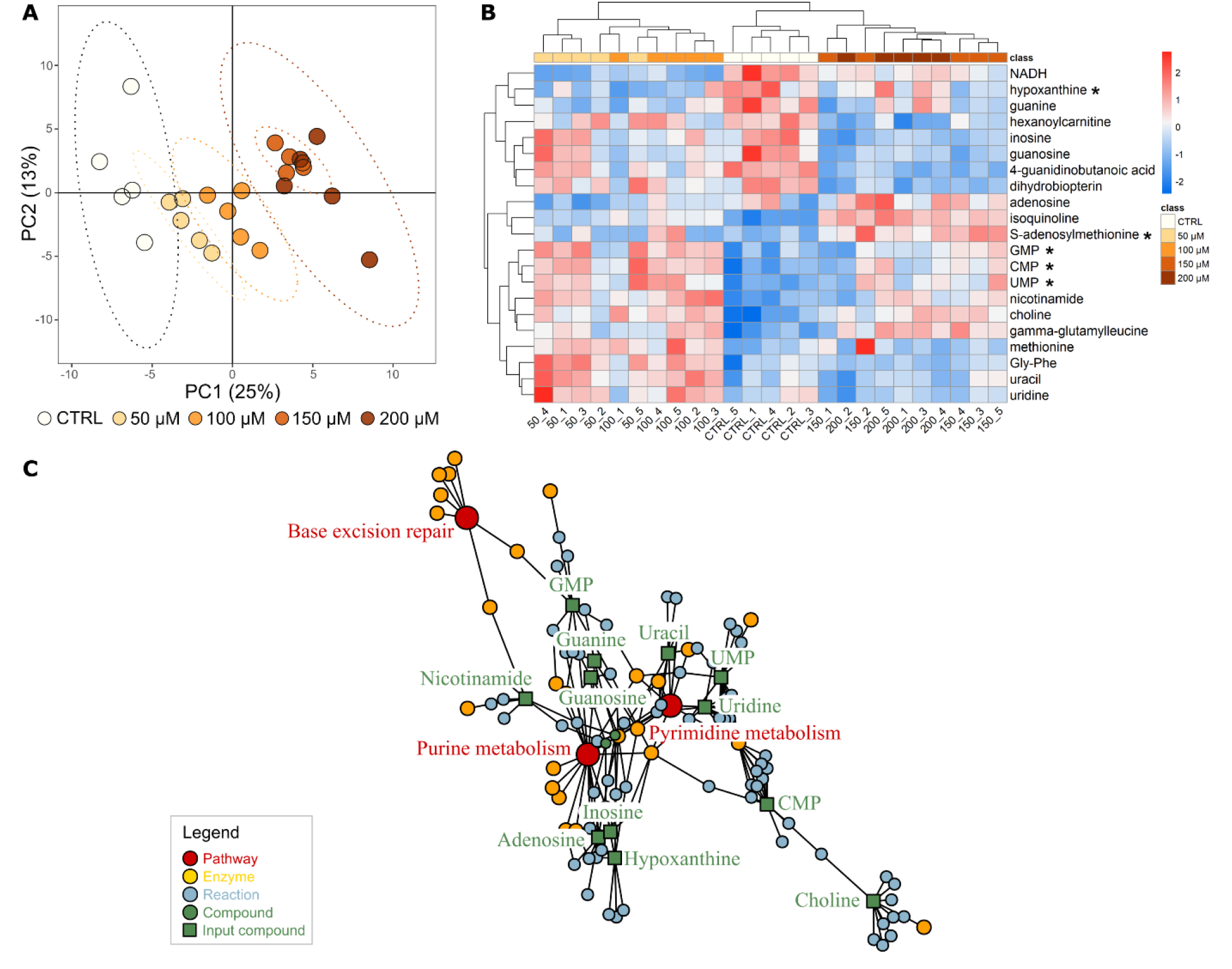
Metabolomic results of the molecular phenotype. **(A)** Unsupervised PCA analysis of metabolomics data. PCA score plot was generated from 2,751 features detected in control and NNK exposed groups after data filtering and normalization. **(B)** Heatmap and hierarchical clustering of identified endogenous metabolites. Each column represents the relative intensity of each metabolite per sample after standard scaling. Asterisks denote metabolites with level 1 identification. **(C)** Network-based pathway enrichment by FELLA of significant endogenous metabolites altered by NNK exposure using the top 150 z-score for diffusion analysis. Red, yellow, blue, and green circles represent the enriched pathways, and potentials enzymes, reactions, and compounds to be also affected. Green squares represent the input metabolites to the enriched pathways.

**Figure 4.**
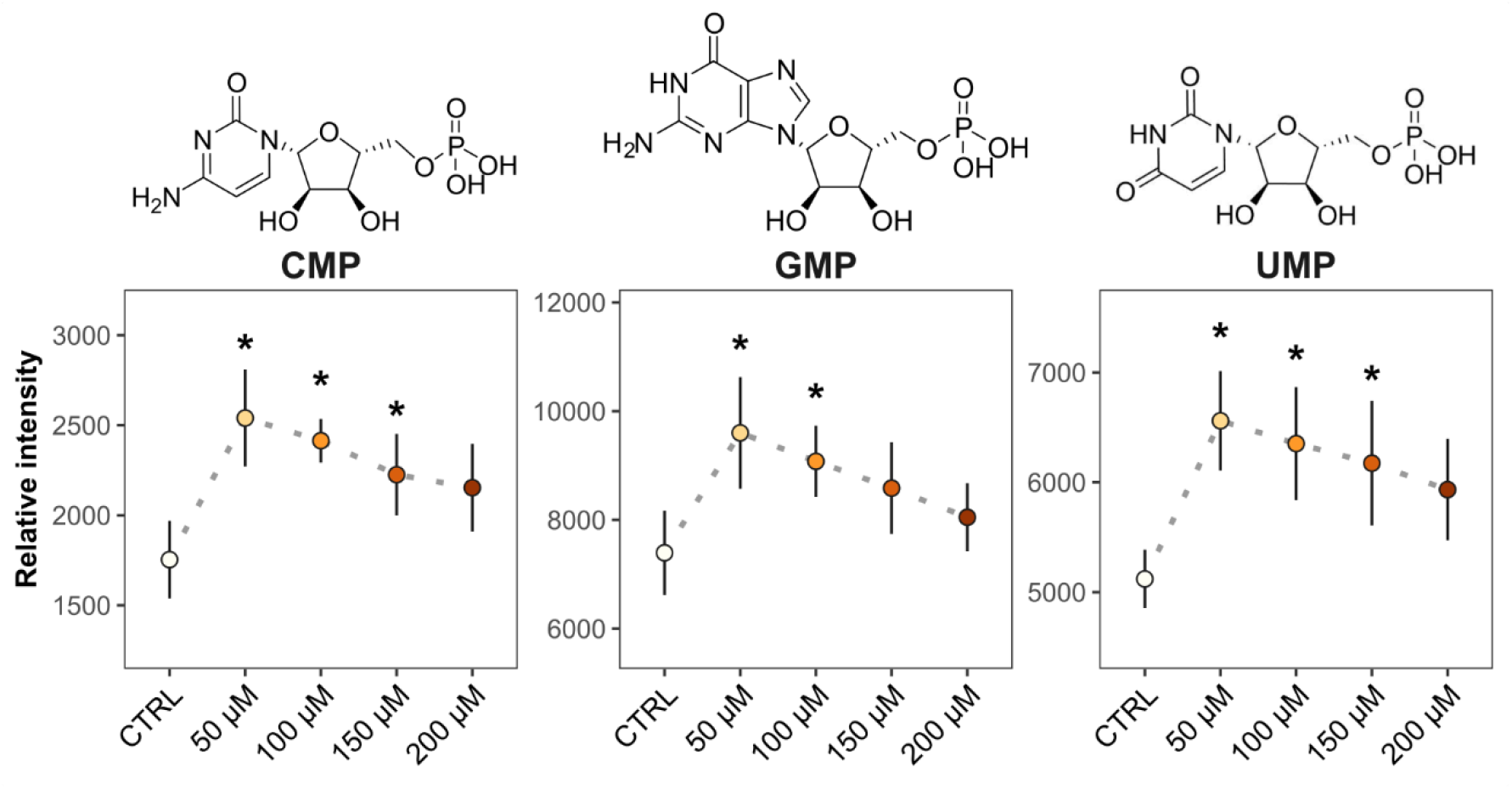
Identified nucleotide monophosphate significantly altered by NNK exposure. Each point represents the mean intensities and error bars the standard deviation. Asterisks denote statistical significance according to multiple group comparison of ANOVA (only significative results of exposed groups vs control are illustrated); * p-value < 0.05. n = 5 samples.

**Table 1.** List of the identified endogenous metabolites altered in zebrafish embryos exposed to NNK, their KEEG ID, FDR adjusted p-value, and identification level (ID level).

| Metabolite | KEEG ID | FDR p-value | ID level |
| --- | --- | --- | --- |
| 4-guanidinobutanoic acid | C01035 | $5.60 \times 10^{-6}$ | 2 |
| adenosine | C00212 | $4.69 \times 10^{-2}$ | 2 |
| choline | C00114 | $3.18 \times 10^{-5}$ | 2 |
| **cytidine monophosphate (CMP)** | **C00055** | **$1.87 \times 10^{-3}$** | **1** |
| dihydrobiopterin | C02953 | $4.18 \times 10^{-4}$ | 2 |
| gamma-glutamylleucine | - | $1.31 \times 10^{-3}$ | 2 |
| glycylphenylalanine (Gly-Phe) | - | $2.28 \times 10^{-4}$ | 2 |
| **guanosine monophosphate (GMP)** | **C00144** | **$1.88 \times 10^{-2}$** | **1** |
| guanine | C00242 | $3.21 \times 10^{-3}$ | 2 |
| guanosine | C00387 | $1.08 \times 10^{-2}$ | 2 |
| hexanoylcarnitine | - | $5.53 \times 10^{-3}$ | 2 |
| **hypoxanthine** | **C00262** | **$3.96 \times 10^{-2}$** | **1** |
| inosine | C00294 | $7.05 \times 10^{-3}$ | 2 |
| isoquinoline | C06323 | $1.60 \times 10^{-10}$ | 2 |
| methionine | C00073 | $5.93 \times 10^{-2}$ | 2 |
| Reduced nicotinamide adenine dinucleotide (NADH) | C00004 | $5.29 \times 10^{-5}$ | 2 |
| nicotinamide | C00153 | $3.64 \times 10^{-3}$ | 2 |
| **S-adenosylmethionine** | **C00019** | **$1.30 \times 10^{-2}$** | **1** |
| **uridine monophosphate (UMP)** | **C00105** | **$6.98 \times 10^{-3}$** | **1** |
| uracil | C00106 | $7.94 \times 10^{-3}$ | 2 |
| uridine | C00299 | $2.39 \times 10^{-2}$ | 2 |

## 4. Conclusions

Our results demonstrate for the first time the impact of NNK exposure to zebrafish embryos. A weak teratogenic and embryotoxic effect was observed with a LOAEL of 200 μM based on lethality. Besides, the detection of NNK, HPBA and NNAL-N-oxide in zebrafish extracts confirmed our zebrafish embryo model to metabolize NNK. We suggest two parallel pathways to be activated: NNK detoxification through NNAL-N-oxide formation and CYP450-mediated α-hydroxylation forming reactive intermediates that cause DNA-adducts eventually driving carcinogenesis. We demonstrated these to be paralleled by activation of BER pathway that counteracts genome instability caused by DNA-lesions occurring upon NNK exposure. Additionally, this is also accompanied by a disruption of purine and pyrimidine metabolisms with nucleotides biosynthesis increasing at the lowest benchmark NNK concentrations. Altogether, these results confirm NNK to be a harmful embryonic agent and demonstrate zebrafish embryos to be a suitable early development model to monitor NNK toxicity reproducing previously observed effects *in-vitro* and in higher order vertebrate models.

## Data availability

Raw mass spectrometry data files (mzXML) an RMarkdown file containing code to reproduce data analysis are available at Zenodo with accession number 4775188

## Author’s contribution

**Carla Merino:** Software, Validation, Formal analysis, Investigation, Data Curation, Writing-Original Draft; Visualization. **Marta Casado:** Methodology, Resources, Investigation. **Benjamí Piña:** Conceptualization, Methodology, Resources, Writing – Review & Editing. **Maria Vinaixa:** Conceptualization, Formal analysis, Writing – Original Draft, Review & Editing, Visualization, Supervision. **Noelia Ramírez:** Conceptualization, Investigation, Supervision, Writing – Review & Editing, Project administration, Funding acquisition.

## Acknowledgements

This research was funded by the Secretaria d’Universitats i Recerca del Departament d’Empresa i Coneixement de la Generalitat de Catalunya through C.M.’s predoctoral grant number 2020 FI_B2 00118 and the Spanish Ministry of Science & Innovation through RTI2018-096175-B-I00 (B.P., M.C.), N.R.’s Juan de la Cierva Incorporación grant No. (IJCI-2015-23158), N.R.’s Miguel Servet contract (CP19/00060) from Instituto de Salud Carlos III, co-financed by Fondo Europeo de Desarrollo Regional (FEDER), Unión Europea, “Una manera de hacer Europa”.

