## Supplementary Table S1 and S2 for "Toxicity of 4-(Methylnitrosamino)-1-(3-pyridyl)-1-butanone (NKK) in early development: a wide-scope metabolomics assay in zebrafish embryos"

---

Maria Vinaixa, PhD
URV, Department of Electronic Engineering
Av. Països Catalans, 26
43007-Tarragona, Spain

**Table S1.** Parameters assessed to classify embryo phenotypes

| Phenotype | Parameters* |
|---|---|
| Death embryos | Non-viable embryos; coagulated embryos; lack of heartbeat; and death embryos/larvae. |
| Severe phenotype | Lack of movement; malformation of eyes, head, mouth, and pectoral fins; modified chorda structure; no-hatched embryos; pericardial and yolk sac oedema; scoliosis; and yolk deformation. |
| Normal phenotype | Hatched embryos without any of the above commented phenotypic parameters. |

* The observation of a single parameter assigned the phenotype

**Table S2.** List of previously reported NNK metabolites (Dator et al., 2018; Hecht, 1998), their abbreviation, theoretical and empirical protonated mass [M+H], empirical retention time ($t_R$), identification level (ID level), and source of LC-MS/MS data.

| Chemical name | Abbreviation | Chemical formula | Theoretical [M+H] | Empirical [M+H] | Empirical $t_R$ (min) | ID level* | MS/MS Refs. |
|---|---|---|---|---|---|---|---|
| 4-Hydroxy-1-(3-pyridyl)-1-butanone | **HPB** | $C_9H_{11}NO_2$ | 166.0863 | 166.0867 | 4.83 | *Not identified* | HMDB |
| 3-Hydroxy-1-(3-pyridyl)-1-butanol | **1,3-Diol** | $C_9H_{13}NO_2$ | 168.1019 | 168.1016 | 1.74 | *Not identified* | (Dator et al., 2018) |
| 4-Hydroxy-1-(3-pyridyl)-1-butanol | **1,4-Diol** | $C_9H_{13}NO_2$ | 168.1019 | 168.1016 | 1.74 | *Not identified* | (Dator et al., 2018) |
| 1,1-Diol | **Unknown 1 (Diol 3)** | $C_9H_{13}NO_2$ | 168.1019 | 168.1016 | 1.74 | *Not identified* | (Dator et al., 2018) |
| 4-Oxo-4-(3-pyridyl)butyric acid | **OPBA** | $C_9H_9NO_3$ | 180.0655 | *Not found* | *Not found* | - | - |
| 4-Hydroxy-4-(3-pyridyl)-butyric acid | **HPBA** | $C_9H_{11}NO_3$ | 182.0812 | 182.0809 | 2 | **1** | STDR |
| 4-(Methylnitrosamino)-1-(3-pyridyl)-1-butanone | **NNK** | $C_{10}H_{13}N_3O_2$ | 208.1081 | 208.1078 | 5.87 | **1** | STDR |
| 4-(Methylnitrosamino)-1-(3-pyridyl)-1-butanol | **NNAL** | $C_{10}H_{15}N_3O_2$ | 210.1237 | 210.1228 | 4.7 | *Not identified* | STDR |
| 4-(Methylnitrosamino)-1-[3-(6-hydroxypyridyl)-1-butanone | **6-OH NNK** | $C_{10}H_{13}N_3O_3$ | 224.103 | *Not found* | *Not found* | - | - |
| 4-[Methyl(nitroso)amino]-1-(1-oxido-3-pyridinyl)-1-butanone | **NNK-*N*-oxide** | $C_{10}H_{13}N_3O_3$ | 224.103 | *Not found* | *Not found* | - | - |
| *N*-(3,4-Dihydroxy-4-(pyridin-3-yl)butyl)-*N*-methylnitrous amide | **γ-OH NNAL (OH-NNAL 2)** | $C_{10}H_{15}N_3O_3$ | 226.1186 | 226.1187 | 5.18 | *Not identified* | (Dator et al., 2018) |
| 4-(Methylnitrosamino)-1-(3-pyridyl-N-oxide)-1-butanol | **NNAL-*N*-oxide** | $C_{10}H_{15}N_3O_3$ | 226.1186 | 226.1187 | 5.18 | **2** | (Dator et al., 2018) |
| - | **Unknown 2 (OH-NNAL 1)** | $C_{10}H_{15}N_3O_3$ | 226.1186 | 226.1187 | 5.18 | *Not identified* | (Dator et al., 2018) |
| 3-(4-(Methyl(nitro)amino)butanoyl)pyridine 1-oxide | **nitro-NK-*N*-oxide** | $C_{10}H_{13}N_3O_4$ | 240.0979 | 240.0981 | 5.18 | *Not identified* | (Dator et al., 2018) |
| 3-(1-Hydroxy-4-(methyl(nitro)amino)butyl)pyridine 1-oxide | **nitro-NAL-*N*-oxide** | $C_{10}H_{15}N_3O_4$ | 242.1135 | *Not found* | *Not found* | - | - |
| - | **Unknown 3** | - | 254.1135 | *Not found* | *Not found* | - | - |
| N-acetylcysteine-PHB 1 | **NAC-PHB 1** | $C_{14}H_{20}N_2O_4S$ | 313.1216 | 313.1212 | 5.22 | *Not identified* | (Dator et al., 2018) |
| N-acetylcysteine-PHB 2 | **NAC-PHB 2** | $C_{14}H_{20}N_2O_4S$ | 313.1216 | 313.1212 | 5.22 | *Not identified* | (Dator et al., 2018) |
| N-acetylcysteine-PHB 3 | **NAC-PHB 3** | $C_{14}H_{20}N_2O_4S$ | 313.1216 | 313.1212 | 5.22 | *Not identified* | (Dator et al., 2018) |

| | | | | | | | |
|---|---|---|---|---|---|---|---|
| 4-Hydroxy-1-(3-pyridyl)-1-butanone glucuronide | **HPB-Gluc** | $C_{15}H_{19}NO_8$ | 342.1183 | *Not found* | *Not found* | - | - |
| 4-(Methylnitrosamino)-1-(3-pyridyl)-1-butanol-O-glucuronide | **NNAL-O-Gluc** | $C_{16}H_{23}N_3O_8$ | 386.1558 | *Not found* | *Not found* | - | - |
| 3,4,5-trihydroxy-6-((nitroso(4-oxo-4-(pyridin-3-yl)butyl)amino)methoxy)tetrahydro-2H-pyran-2-carboxylic acid | **α-OH-methyl-NNK-Gluc** | $C_{16}H_{21}N_3O_9$ | 400.1351 | *Not found* | *Not found* | - | - |
| 3,4,5-Trihydroxy-6-(((4-hydroxy-4-(pyridin-3-yl)butyl)(nitroso)amino)methoxy)tetrahydro-2H-pyran-2-carboxylic acid | **α-OH-methyl-NNAL-Gluc** | $C_{16}H_{23}N_3O_9$ | 402.1507 | 402.1492 | 5.03 | *Not identified* | (Dator et al., 2018) |

***** Identification level according to Schymanski et al. (Schymanski et al., 2014). Identification was performed by matching the empirical MS/MS spectra with the MS/MS of NNK metabolites pure chemical standards (STDR) (Level 1). When pure chemical standards were not available, annotation was performed by comparing the empirical MS/MS spectra with the MS/MS spectra available in the HMDB (Wishart et al., 2018) or in the literature (Dator et al., 2018) (level 2). *Not found*: NNK metabolites are those whose empirical protonated mass was not within 5 ppm mass error. *Not identified*: NNK metabolites are those whose empirical MS/MS spectra did not match with the MS/MS spectra of NNK metabolites pure chemical standards, databases, or literature. *Unknow*: compounds are novel NNK metabolites described by Dator et.al. (Dator et al., 2018).
